# Secondary evolve and re-sequencing: an experimental confirmation of putative selection targets without phenotyping

**DOI:** 10.1101/722488

**Authors:** Claire Burny, Viola Nolte, Pierre Nouhaud, Marlies Dolezal, Christian Schlötterer

**Affiliations:** Institut für Populationsgenetik, Vetmeduni Vienna, Vienna, Austria; Vienna Graduate school of Population Genetics, Vetmeduni Vienna, Vienna, Austria

**Keywords:** experimental evolution, *Drosophila simulans*, repeatability of evolution, evolve and re-sequence

## Abstract

Evolve and re-sequencing (E&R) studies investigate the genomic responses of adaptation during experimental evolution. Because replicate populations evolve in the same controlled environment, consistent responses to selection across replicates are frequently used to identify reliable candidate regions that underlie adaptation to a new environment. However, recent work demonstrated that selection signatures can be restricted to one or a few replicate(s) only. These selection signatures frequently have a weak statistical support, and given the difficulties of functional validation, additional evidence is needed before considering them as candidates for functional analysis. Here, we introduce an experimental procedure to validate candidate loci with weak or replicate-specific selection signature(s). Crossing an evolved population from a primary E&R experiment to the ancestral founder population reduces the frequency of candidate alleles that have reached a high frequency. We hypothesize that genuine selection targets will experience a repeatable frequency increase after the mixing with the ancestral founders if they are exposed to the same environment (secondary E&R experiment). Using this approach, we successfully validate two overlapping selection targets, which showed a mutually exclusive selection signature in a primary E&R experiment of *Drosophila simulans* adapting to a novel temperature regime. We conclude that secondary E&R experiments provide a reliable confirmation of selection signatures that are either not replicated or show only a low statistical significance in a primary E&R experiment. Such experiments are particularly helpful to prioritize candidate loci for time-consuming functional follow-up investigations.

## INTRODUCTION

Experimental evolution provides the opportunity to study evolutionary processes over time scales short enough to be followed experimentally (Garland and Rose 2009; Kawecki et al. 2012). The combination of high-throughput sequencing with experimental evolution (Evolve and Re-sequence, E&R) has been widely used to identify adaptive alleles across multiple replicates starting from the same reservoir of standing variation in highly similar, well-controlled environments (Turner et al. 2011; Long et al. 2015; Schlötterer et al. 2015). E&R studies successfully characterized the genomic responses during adaptation to novel selective pressures usually on organisms with short generation times (e.g.: Turner and Miller 2012; Burke et al. 2014; Lenski 2017; Papkou et al. 2019; Remigi et al. 2019). Laboratory natural selection experiments using the E&R framework studied responses to thermal (Orozco-terWengel et al. 2012; Tobler et al. 2014; Michalak et al. 2019) or desiccation stress (Schou et al. 2014), starvation (Michalak et al. 2019) and salt- and cadmium-enriched environments (Huang et al, 2014). The advantage of E&R studies starting from natural variation is that adaptation is possible without de novo mutations (Teotónio et al. 2009). Hence, even organisms with moderate experimental population sizes, such as *Drosophila*, are able to adapt to novel conditions within experimentally feasible time scales. Furthermore, when the starting variation is sampled from a natural population, E&R studies provide direct information about the frequency of the selected alleles in the wild (Barghi et al. 2019).

Standard statistical tests applied to E&R data (e. g. Cochran Mantel Haenszel (CMH) test (Agresti, 2002; Spitzer et al, unpublished data) or Generalized Linear Modeling (Phillips et al. 2018)) require parallel selection responses across replicates. Two different, not mutually exclusive, factors can severely compromise the detection of selection targets based on these approaches. Polygenic adaptation to a new trait optimum results in reduced genomic parallelism across replicates (Franssen et al. 2017; Barghi et al. 2019). Furthermore, selected alleles with low starting frequencies are not only less likely to reach a detectable selection signature, but genetic drift, i.e. chance, also results in lower repeatability across replicates (Lenormand et al. 2016). One further complication for the identification of selection targets with low starting frequencies arises from hitchhiking of SNPs shared with haplotypes carrying the favorable allele (Nuzhdin and Turner 2013; Tobler et al. 2014; Franssen et al. 2015). In this case, the limited number of recombination events during the experiment results in large genomic regions with selection signatures when selection operates on low frequency alleles, that make the identification of individual candidate genes impossible.

The functional characterization of selected alleles in E&R studies is an important next step for a better understanding of adaptation processes, but despite the recent advances based on the CRISPR/Cas9 technology (Bassett et al. 2013), the functional characterization of different alleles in a standardized genetic background is still a challenging and time-consuming task. This implies that investigators are well-advised to have high confidence in alleles that are going to be functionally tested.

We propose a simple experimental procedure to validate candidate regions with weak statistical support, either due to a weak selection signature across replicates or replicate-specific selection signatures. The basic idea of this approach is that an evolved population is “diluted” with ancestral genotypes. This reduces the frequency of putatively selected alleles and the reproducible increase in frequency of selected alleles in multiple replicates evolving under the same selection regime (secondary E&R) serves as a validation of candidate regions. Because secondary E&R experiments provide the opportunity for additional recombination events, we also evaluated whether this approach increases the mapping resolution, which is particularly important for low frequency beneficial alleles.

Applying secondary E&R to a candidate region identified in *D. simulans* populations that have been exposed to a novel constant hot environment at 23°C for 70 generations, we demonstrate that candidate selection targets can be experimentally confirmed.

## NEW APPROACHES

Previously, experimental evolution studies exposed laboratory evolved populations to selection regimes in the opposite direction (reverse evolution) (Teotónio and Rose 2001; Porter and Crandall 2003; Teotónio et al. 2009). The secondary E&R design introduced here, also relies on already laboratory selected populations, but rather than changing the selection regime, the same selection regime is applied after manipulating the evolved population. Secondary E&R is designed to provide researchers additional confidence about selection targets by repeating a selection signature in replicate populations after adding genotypes from the founder population, which reduces the frequency of selected alleles. The repeated, parallel frequency increase of candidate regions provides a reliable confirmation of selection targets that were either having a weak selection signal or were only detected in a single replicate.

## RESULTS

### Discovery of candidate SNPs: primary E&R

Three replicates of a *D. simulans* founder population were maintained in a constant hot environment (23°C) for 70 non-overlapping generations. Sequencing pools of 1,250 individuals (Pool-Seq, Kofler and Schlötterer 2013; Schlötterer et al. 2014; Table SI 1) resulted in a catalogue of 2,560,538 polymorphic SNPs (see Methods, Table SI 2). We identified candidate SNPs by contrasting allele frequency changes (AFC) between ancestral and evolved populations with a CMH test after accounting for drift using a 1% empirical FDR threshold (see Methods). Since p-values obtained from contingency tables tests are affected by coverage, we also accounted for coverage heterogeneity among samples (56x – 261x, Table SI 3) by weighting p-values following the Iterative Hypothesis Weighting procedure (IHW, Ignatiadis et al. 2016) (see Methods). The genome-wide analysis identified a candidate region of 1.628Mb on chromosome arm 3R with a pronounced AFC between ancestral and evolved populations (Fig. 1., top left, the full genomic analysis will be published elsewhere).

**Figure 1.**
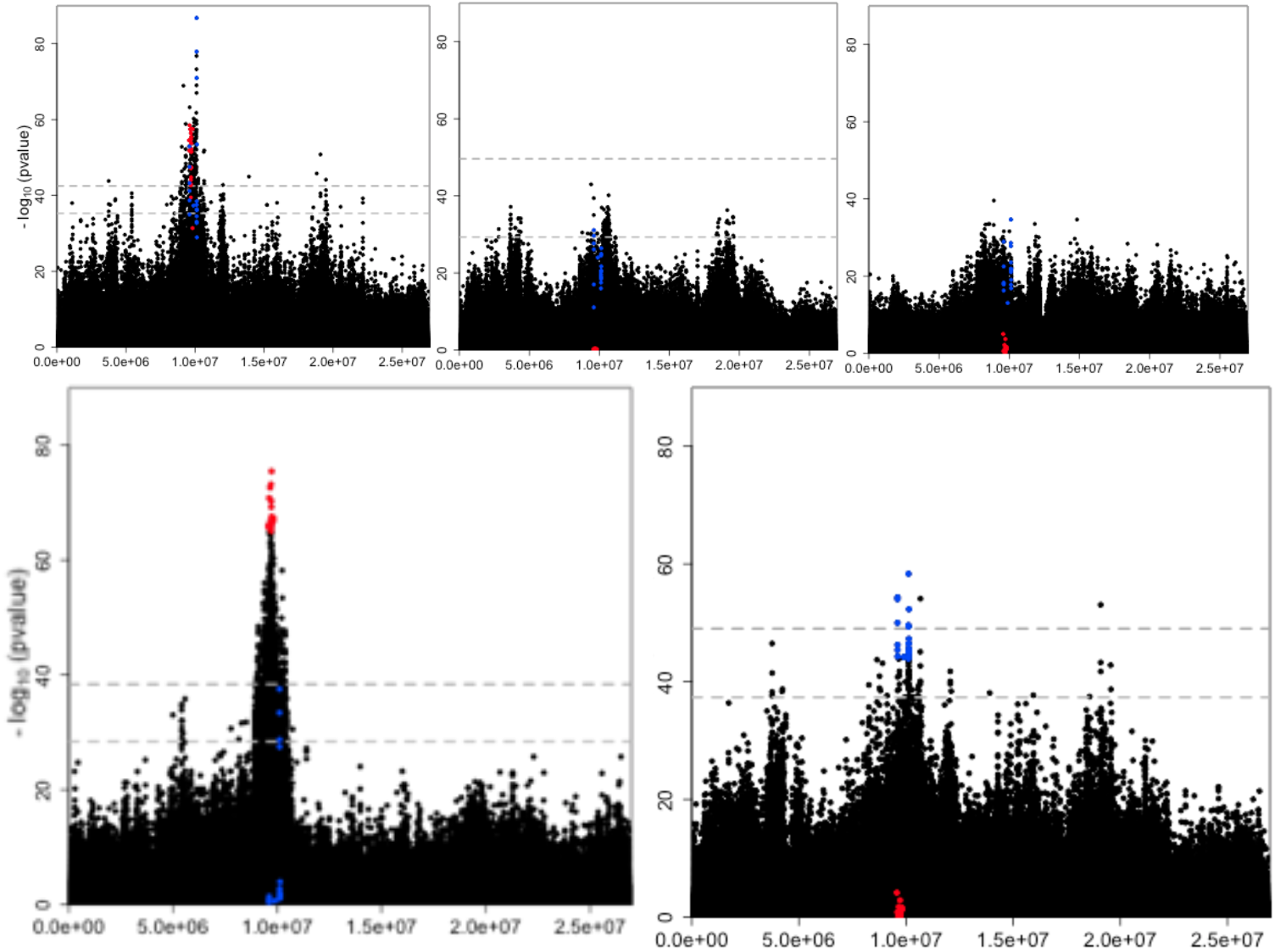
Replicate-specific selection signatures in the primary E&R study. Manhattan plots of chromosome arm 3R displaying the negative log10-transformed weighted p-values of 680,937 SNPs for different statistical tests. A) CMH_x,y,z_ (175/443), B) FET_y_ (0/122), C) FETz (0/0), D) FET_x_ (660/1,776) and E) CMH_y,z_ (9/85). The number of candidates at 1%/5% empirical FDR thresholds for each test are given in parenthesis. The gray dotted line shows the 1% (upper) and 5% (lower) empirical FDR thresholds of the corresponding test, computed over the autosomes from neutral simulations assuming no linkage. At the 1% empirical FDR threshold, CMH_x,y,z_ and FET_x_ identify a candidate peak region of 169 (9,042,023-10,670,451bp, 1.628Mbp) and 660 (9,000,008-10,384,933bp, 1.385Mbp) SNPs. The overlap between these two tests is 92 significant SNPs spanning 1.343Mb (see SI Fig. 1. for a close up of this genomic region). In all panels the top 20 SNPs from FET_x_ and CMH_y,z_ are highlighted in red and in blue.

The power of the CMH test relies on the experimental replicates to detect putative targets of selection. However, its power is limited when candidates are not shared across replicates. Analyzing this genomic region separately for each of the replicates using a Fisher’s Exact Test (FET) indicated considerable heterogeneity among them: among the SNPs with the most significant CMH p-values across all three replicates, the top 20 SNPs in the FET of replicate x were only significant in replicate x (FET_x_, Fig. 1., bottom left, top center, top right, red), with 16 SNPs being close to fixation. Removing replicate x from the CMH analysis and using only replicates y and z, we obtained a much weaker selection signature in the CMH test (CMHy,z, Fig. 1., bottom right). Only three of the 20 most significant SNPs of this analysis (CMH_y,z_) were overlapping with the most significant SNPs of the analysis including x (CMHx,_y,z_). Instead, the 20 most significant SNPs of CMH_y,z_ changed in both replicates y and z with a mean AFC of 0.55. This AFC is less pronounced than the one observed for the significant SNPs of replicate x (0.96). This heterogeneity among replicates suggested that at least two distinct classes of haplotypes were selected.

We further scrutinized the hypothesis of at least two distinct selected haplotypes and plotted the AFC of the two sets of top 20 SNPs in the candidate region on chromosome arm 3R (Fig. 3.): 20 SNPs from FET_x_ and 20 SNPs from the joint analysis of replicates y and z, i.e. CMH_y,z_. The two sets of candidate SNPs displayed group-specific AFC; one set showed a pronounced AFC in replicate x and the other one in replicate z, but almost no change in the other (Fig. 3., Fig. SI 1.).

### Validation of candidate SNPs: secondary E&R

The primary E&R study provided two sets of candidate SNPs. One set of candidates increased strongly in replicate x only, while the other set of candidates increased weakly in the two replicates y and z. To demonstrate that both sets of SNPs are associated with a selection target, we aimed to validate both selection signatures experimentally. Reasoning that fewer replicates are needed to confirm strong selection, only two diluted replicates were generated from evolved replicate x (x.1 and x.2), while three diluted replicates were generated from evolved replicate z (z.1, z.2 and z.3) which showed the weakest response in the initial E&R experiment. For both secondary E&R experiments we added flies from a reconstituted founder population (Nouhaud et al. 2016) aiming for a starting frequency around 0.5 for the most prominent candidate SNPs (see Fig. SI 2). This starting frequency of the candidate SNPs in the secondary E&R ensures a deterministic selection response and still provides sufficient opportunity for frequency increase.

After 30 generations of evolution at the same culture conditions, we sequenced the founders (D0) and evolved replicates (D30) of the secondary E&R experiments (see Fig. 2. for an overview). We contrasted the dynamics of the two groups of top candidate SNPs in each of the replicates in the primary and secondary E&R experiments over four time points (F0, F70, D0, D30). A very pronounced frequency increase can be noted in both the primary and secondary E&R experiments in the focal replicate from which the candidates were obtained (Fig. 3., Fig. SI 3). From an average starting allele frequency of 0.52 and 0.31 the candidate SNPs reach a mean final frequency of 0.98 (x) and 0.73 (z) in the replicates of the secondary E&R. The consistent AFC in the primary and secondary E&R experiments confirms a high repeatability of selection. Also, the candidate SNPs from the non-focal replicate consistently failed to show selection signatures (Fig. 3., Fig. SI 3). The only exception are 4 SNPs from the candidate set of replicate z, which also increased in frequency in the primary and secondary E&R of replicate x (Fig. 3., Fig. SI 3, Fi. SI 6). Because the AFC was less pronounced than the one of the focal candidate SNPs of replicate x, we conclude that these SNPs may be shared between the two alternatively selected haplotype classes.

**Figure 2.**
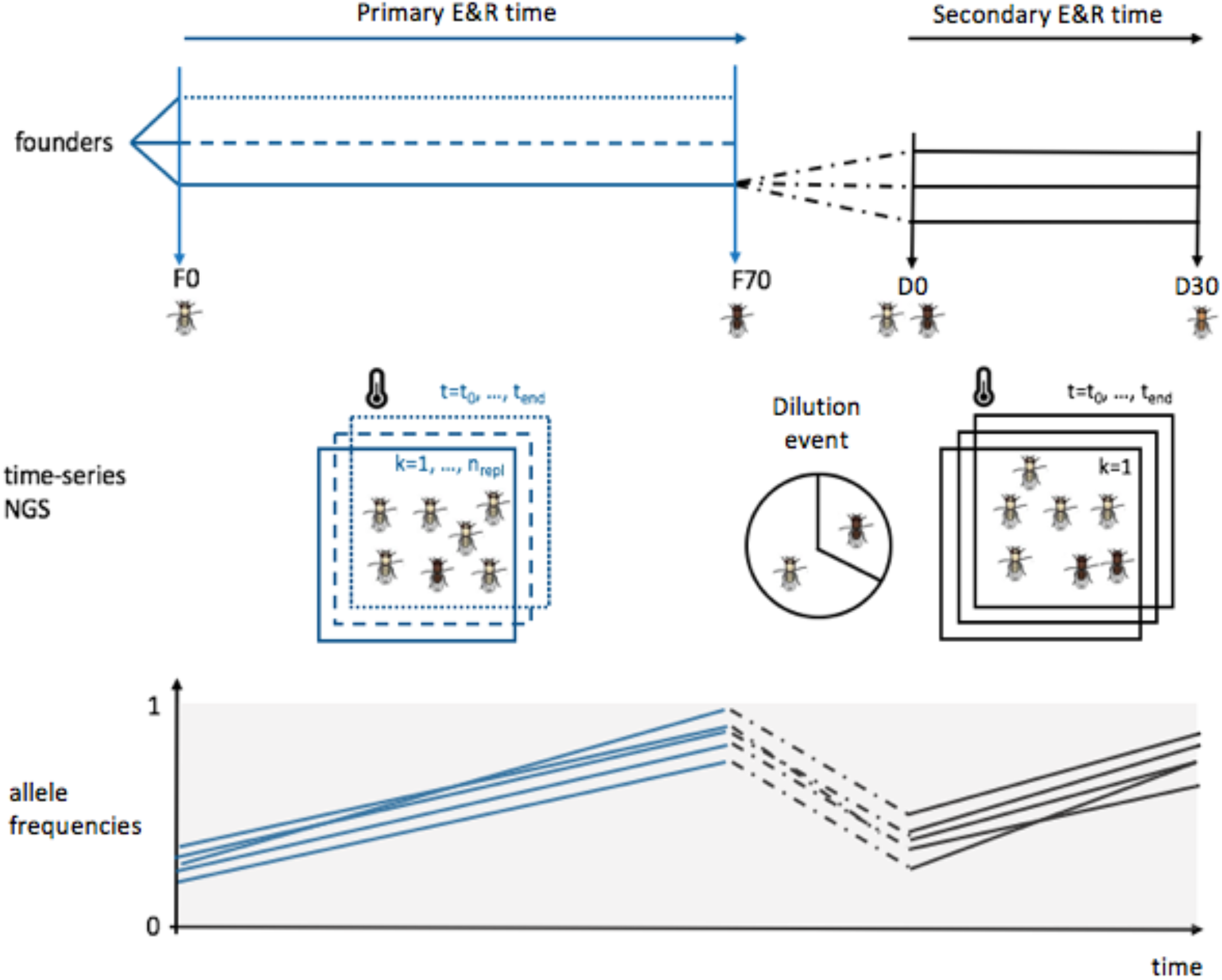
Schematic outline of the experimental design. Three replicated populations of flies starting from the same founders evolved in parallel during 70 generations (primary E&R, indicated in blue). The darkness of the flies symbolizes the level of adaptation to the new environment. Fora given evolved replicate, the evolved flies are “diluted” with ancestral genotypes and independent replicates evolving for an additional 30 generations under the identical environmental conditions as in the primary E&R (secondary E&R, indicated in black). The bottom panel indicates the allele frequency changes of candidate SNPs during the experiments. In the primary E&R the allele frequency increases (blue). By adding ancestral genotypes, the frequency of the candidate SNPs is decreased (black dotted lines). 30 generations of the secondary E&R result in a repeated frequency increase of the candidate SNPs, confirming non-neutral evolution.

**Figure 3.**
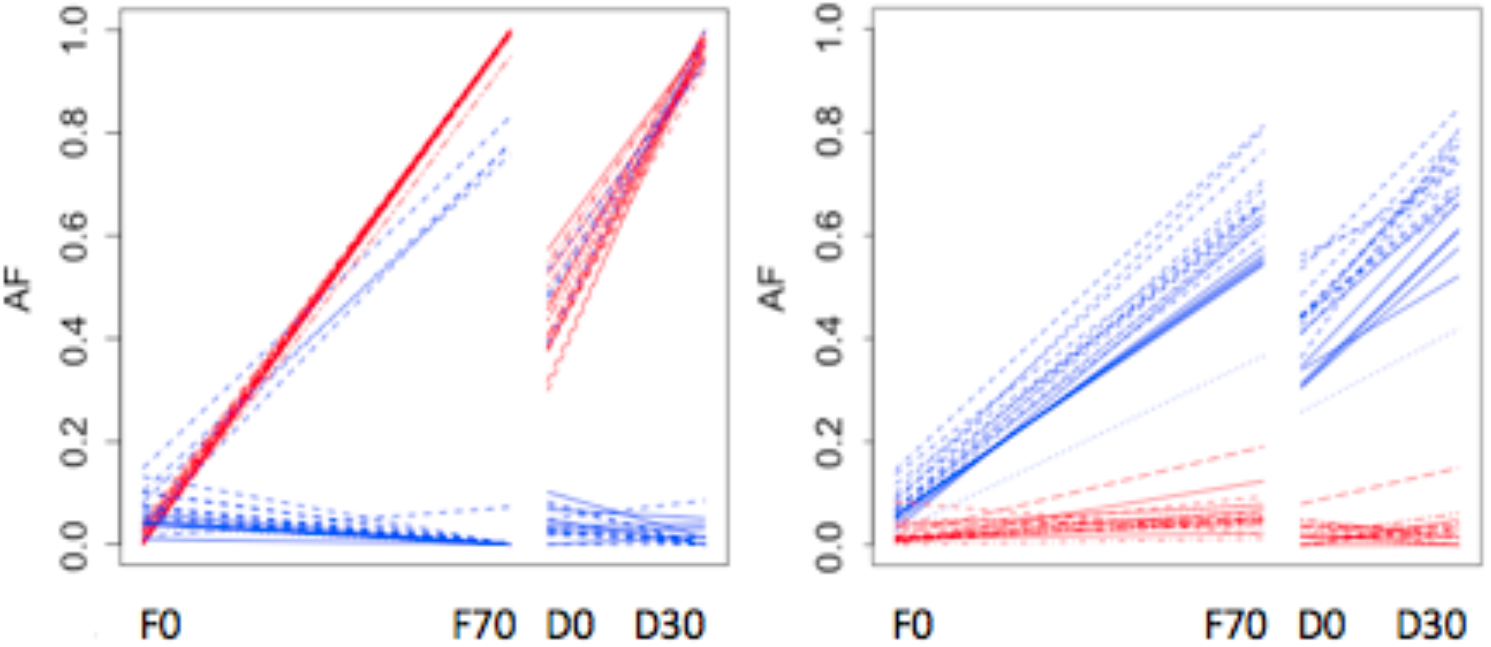
Allele frequency changes of the 20 most significant SNPs from FET_x_ (red) and CMH_y,z_ (blue) for the primary E&R (generation F0-F70) and secondary E&R (D0-D30). The left panel shows experiment x and the right panel experiment z. Only first replicate the secondary E&R is shown for each experiment, for the other replicates, see supplement.

For a more complete picture we expanded our analysis of the 20 most significant SNPs to all significant SNPs (FDR<0.01) of the primary E&R. We jointly plotted the distribution of selection coefficients obtained from the primary and secondary E&R experiments (see Methods). Consistent with the previous analyses, all candidate SNPs had a selection coefficient larger than zero in their focal replicate - independently of whether primary or secondary E&R experiments were analyzed (Fig. 4.a., Fig. SI 4). The inferred selection coefficients for replicate x are about twice as high as the ones for replicate y. The mean selection coefficients from the 20 candidate SNPs are 0.26 and 0.27 for diluted replicates from x (0.26 in the primary E&R) and 0.08, 0.09, 0.12 for diluted replicates from z (0.09 in the primary E&R). As expected the selection coefficients of the non-focal top 20 candidate SNPs were distributed around zero.

**Figure 4.**
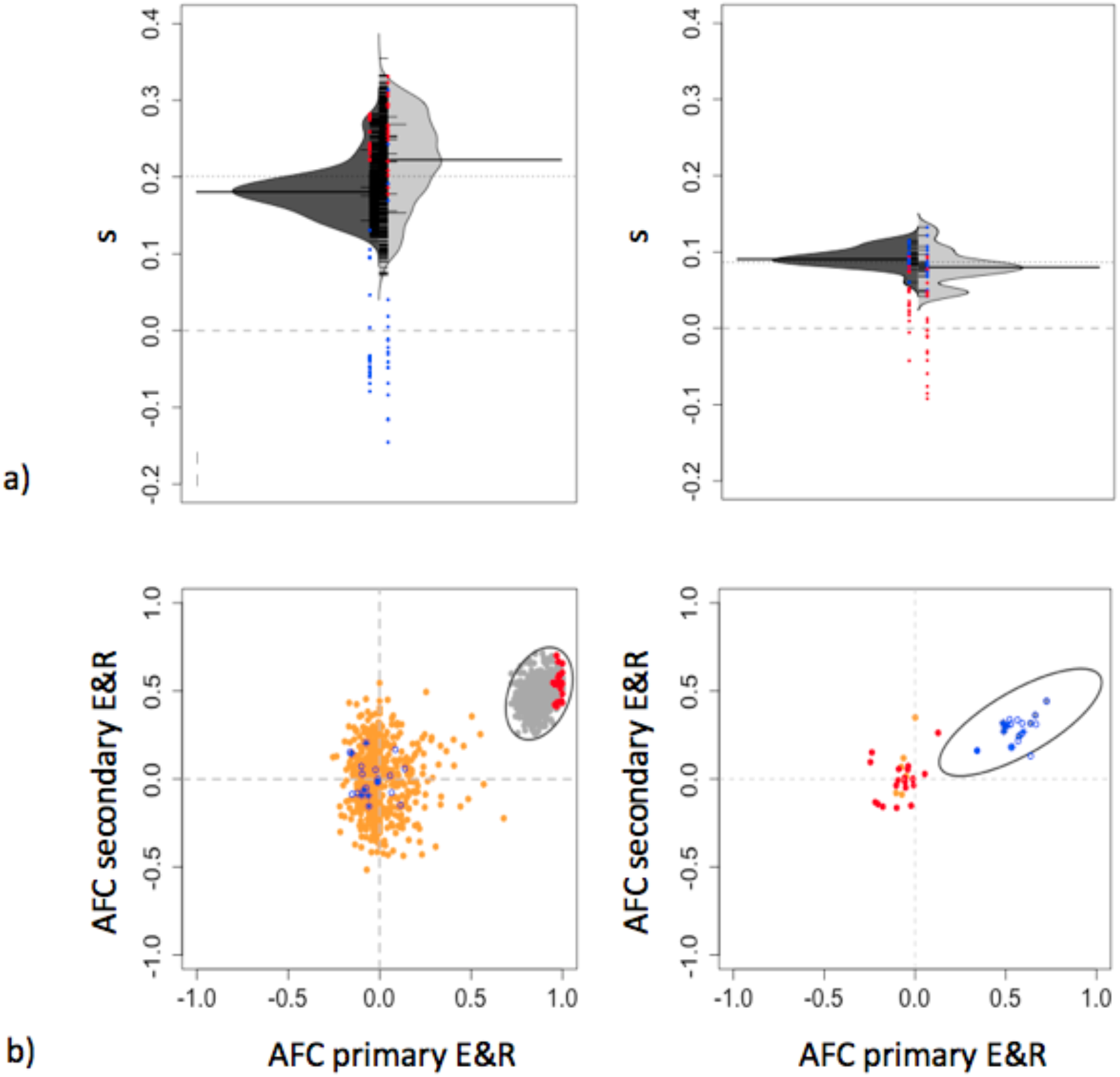
Repeatability of selection signatures in primary and secondary E&R. A) Selection coefficients are very similar. Symmetrical violin plots of the selection coefficients from primary E&R (dark gray) and the first replicate of the secondary E&R experiment (light gray) for candidates in the region of interest. Black segments represent the median per sample. Ticks indicate SNPs. Left: experiment x. Right: experiment z. The 20 most significant SNPs from FET_x_ (red) and CMH_y,z_ (blue) are shown in color. B) Parallel changes in allele frequencies. Observed allele frequency changes for candidate SNPs (empirical FDR <1%) in replicate x (left) are shown in gray. For comparison, the expected neutral allele frequency changes based on the same starting frequency and coverage and a single simulation run are shown in orange. The 20 most significant SNPs from FET_x_ and CMH_y,z_ are shown in red and blue. Since, for replicate z no SNP exceeded the empirical FDR of 1% in the primary E&R, only the top 20 SNPs are shown (right panel). Ellipses around the empirical focal SNPs indicate the 99% probability range to visualize the bivariate densities.

Finally, to evaluate the influence of genetic drift, we simulated the dynamics of the significant SNPs (FDR<0.01) in the primary E&R under neutrality and compared them to their observed dynamics (Fig. 4.b., Fig. SI 5 and Methods). Plotting the pairwise observed and simulated neutral AFC of the primary E&R against the AFC of the secondary E&R experiment, we find that the simulated data are clearly distinct from the experimental ones. The significant SNPs of the experimental data cluster together in the upper right quadrant and do not overlap with neutral simulations, showing that genetic drift cannot explain the concordant signatures of the significant SNPs. As expected the separation of neutral and selected SNPs was clearer for the replicate x, where selection was stronger (Fig. 4.a.).

### No increased mapping resolution for the selection target

Given that the dilution reduced the frequency of the selection target, we anticipated that additional recombination events occurring during the repeated spread of the selection targets would also increase the mapping resolution. Nevertheless, we noted that the selection signature was broader in the secondary E&R experiment than in the primary one (Fig. SI 6). Hence, despite the highly repeatable selection signature of candidate SNPs, the secondary E&R experiment did not yield more confidence about the focal target of selection than the primary E&R experiment.

## DISCUSSION

One of the undisputed advantages of experimental evolution is that the precise experimental conditions are known, which allows to impose the same selection pressure on different populations and time points in a replicated manner. Hence, unless strong epistatic interactions dominate, it should be possible to confirm selected variants by experimentally manipulating allele frequencies in the population in which a favorable variant spread.

In this report we introduce a simple manipulation of the evolved populations. By adding unevolved genotypes, we reduce the frequency of the selection target, which provides the opportunity to monitor a repeatable frequency increase of selected alleles in replicated populations. Our results demonstrate that this novel approach accurately recovers the selection signature of candidate SNPs. Despite the mapping resolution of the primary E&R experiment could not be improved, it is striking how consistent the selection coefficients of the top candidate SNPs were replicated in the secondary E&R experiments, in particular because no phenotypes were measured and the actual selective force is not yet characterized.

We propose that secondary E&R experiments with unevolved genotypes provide an attractive approach to experimentally validate selection signatures. This is particularly important for either non-replicated or small allele frequency changes - both signatures of polygenic adaptation. The power of secondary E&R experiments is well-illustrated in our proof of principle study, in which no single SNP passed the genome-wide significance threshold in this genomic region in the primary E&R experiment in replicate z. Only by combining two replicates, y and z, we identified significant candidates, which could be confirmed in the secondary E&R experiment. Thus, we demonstrated that even populations with weak selection signatures can be used to confirm the presence of selection, which could not be recognized before.

Secondary E&R experiments are not fast, the 30 generations of this experiment took about 14 months, but the maintenance of replicate populations does not require many resources and provides therefore a very good approach to experimentally validate genomic regions experiencing selection. Mapping of causative variants could not be achieved in this pilot study and requires alternative approaches to do so. Nevertheless, the dynamics of selected genomic regions are highly informative of the underlying genetic architecture of beneficial mutations. Polygenic adaptation to a novel trait optimum displays characteristic dynamics (Franssen et al, 2017), which are best detected in multiple replicates. We anticipate that the analysis of multiple replicates in secondary E&R experiments will provide an unprecedented opportunity to study replicated dynamics of selection targets in order to understand the architecture of adaptation. It is also conceivable to use this experimental setup to study the dynamics of a given selected region in an alternative selection regime.

A particularly interesting pattern could be confirmed in this study: two different haplotype classes are carrying adaptive variants that increase fitness of the populations in a novel hot environment. It is particularly remarkable that the two groups of haplotypes seem to be mutually exclusive - we see either one or the other increasing in frequency in the primary E&R experiment. Also in the secondary E&R experiments we see no evidence of parallel selection of both haplotype classes, but their different starting frequencies in the secondary E&R considerably decrease the opportunity for a strong frequency increase of the haplotype with the lower starting frequency. The mapping resolution is not high enough to determine whether the same gene is carrying a beneficial mutation in both haplotype classes or different genes are selected. Thus, similar to many other E&R studies, a good strategy for fine mapping is needed to answer these questions.

## MATERIALS AND METHODS

### The Primary E&R Experiment

#### Experimental Population and Selection Regime

We collected a natural *D. simulans* population 10 km North of Stellenbosch, South Africa, in February and March 2013 and established isofemale lines that were maintained in the laboratory for approximately eight generations. For starting the primary E&R experiment, three mated females from each of 426 isofemale lines were combined three times to generate three replicates of the ancestral population (replicates x, y and z) in F0. They were subsequently maintained as independent populations with a census population size of 1,250 and non-overlapping generations under a constant 23°C temperature regime with a 12 hour light/12 hour dark cycle (LD 12:12) for 70 generations (F70). The 426 lines used for constituting the ancestral population were maintained as isofemale lines.

#### Creation of a Bona Fide SNP Catalogue for the Primary E&R study

We generated Pool-Seq data for the 3 replicates of F0 from females only and for the 3 replicates in F70 (sex ratio ~ 50:50). DNA extraction, barcoded library preparation and sequencing followed standard procedures and are given in Supplementary Table I. We followed standard approaches for quality control, read mapping, read filtering, trimming as well as SNP calling and SNP filtering.

We used libraries with different insert sizes, which can result in false positives (Kofler et al, 2016). To account for this, we expanded the double-mapping procedure suggested by Kofler et al, 2016, and used three different mappers (NovoAlign (http://novocraft.com), Bowtie2 (Langmead and Salzberg, 2012) and BWA-MEM (Li and Durbin, 2009)). We filtered for biallelic SNPs outside of repeat regions, and removed SNPs from positions outside the 99% quantile in terms of genome wide coverage. From this set of pre-filtered SNPs we keep only those for which the SNP frequency did not differ between all three mappers (p>0.01, after FDR correction). We call this procedure triple-mapping. This resulted in a set of 2,560,538 high quality SNPs. Details are given in Supplementary Material I.

#### Identifying Regions under Selection in the Primary E&R

We performed Fisher’s exact tests (FET) between the ancestral F0 and the evolved F70 generation within each replicate and Cochran-Mantel-Haenszel tests (CMH) (Agresti, 2002) across replicates. As coverage variability (see Supplementary Table II) affects the power of FET and CMH tests, we used the independent hypothesis weighting (IHW) procedure (Ignatiadis et al, 2016) to weight the empirical p-values using the mean coverage at each SNP calculated from all replicates included in any particular test, as a covariate.

To determine the list of candidate SNPs, we ran neutral forward Wright-Fisher simulations for each replicate based on *N_e_* estimates (Table 1) that we obtained for autosomes and the X chromosome using the poolSeq package (Taus et al, 2017). Neutral p-values were also submitted to the IHW procedure. Candidate SNPs were declared at a 1% FDR cut off, applying a conservative nonparametric empirical FDR estimator (Strimmer, 2008) using the weighted p-values from our simulations and the weighted p-values from our observed data. This was done separately for FET and CMH tests for each replicate.

**Table 1.** Autosomal *N_e_* estimates of the primary and secondary E&R experiments.

|  | Replicate x | Replicate y | Replicate z |
| --- | --- | --- | --- |
| Primary E&R | 206 | 263 | 226 |
| Secondary E&R | 134, 144 | - | 216, 193, 167 |

Selection coefficients were determined for each SNP in each replicate on pseudo-count data (detailed in Supplementary Material I) using the poolSeq package assuming a dominance coefficient of 0.5.

### The Secondary E&R Experiment

#### Experimental Population, Selection Regime and Sequencing

Based on the primary E&R selection signature screen, we picked a candidate region on 3R (region details in Supplementary Figure 1) for further investigation. This region showed a very strong signal of positive selection in a CMH test across replicates x, y and z. We used evolved flies from replicates x and z after 77 generations of evolution in the primary E&R experiment (F77) to set up a secondary E&R experiment in which the evolved flies were mixed with flies from a reconstituted ancestral population (Nouhaud et al, 2016, Supplementary Figure 2). We call this generation D0. Selection targets are expected to increase in frequency again in the secondary E&R experiment, which used the same culturing conditions as the primary E&R experiment.

Mixing proportions of ancestral and evolved populations to create D0 were chosen such that selected SNPs in our candidate region had allele frequencies of approximately 0.5 in D0: for replicate x, a 30:70 ratio between evolved and reconstituted ancestral flies, and for replicate z, a 50:50 ratio, respectively. We created two replicates for D0 for replicate x (x.1 and x.2), and three replicates for the diluted replicate z (z.1, z.2 and z.3). Replicates for D0 and D30 were subjected to Pool-Seq.

#### Validation of Signatures of Selection in the Secondary E&R

Selection coefficients and neutrality tests were performed exactly as described for the primary E&R experiment.

## ACKNOWLEDGMENTS

We thank all members of the Institute of Population Genetics for feedback and support, especially Neda Barghi, Rui Borges, Lukas Endler, Andreas Futschik, Anna Maria Langmüller, Sonja Lečić, Kathrin Otte. Thomas Taus first proposed the secondary E&R. This work was supported by the Austrian Science Fund (FWF, grant W1225) and an European Research Council (ERC) grant (ArchAdapt).

## Data availability

Raw Pool-Seq data will be uploaded to SRA XXX and available upon publication.

## AUTHORS CONTRIBUTION

V.N. performed experiments. P.N. did the first preliminary analysis of the primary E&R. C.B. analyzed the data. M.D. provided statistical support. C.B., C.S. wrote the paper.

## REFERENCES

Agresti A. 2002. Categorical Data Analysis. John Wiley & Sons, 2nd Edition.

Barghi N, Tobler R, Nolte V, Jakšćc AM, Mallard F, Otte KA, Dolezal M, Taus T, Kofler R, Schlötterer C. 2019. Genetic redundancy fuels polygenic adaptation in *Drosophila*. PLoS Biol 17(2):e3000128.

Bassett A, Tibbit C, Ponting CP, Liu J. 2013. Highly efficient targeted mutagenesis of *Drosophila* with the CRISPR/Cas9 system. Cell Rep 4(1):220–228.

Blount ZD, Lenski RE, Losos JB. 2018. Contingency and determinism in evolution: Replaying life’s tape. Science 362(6415):eaam5979.

Burke MK, Liti G, Long AD. 2014. Standing genetic variation drives repeatable experimental evolution in outcrossing populations of Saccharomyces cerevisiae. Mol Biol Evol 31(12): 3228–3239.

Fragata I, Simões P, Lopes-Cunha M, Lima M, Kellen B, Bárbaro M, Santos J, Rose MR, Santos M, Matos M. 2014. Laboratory selection quickly erases historical differentiation. PLoS One 9(5):e96227.

Franssen S, Kofler R, Schlötterer C. 2017. Uncovering the genetic signature of quantitative trait evolution with replicated time series data. Heredity 118(1):42.

Franssen S, Nolte V, Tobler R, Schlötterer C. 2015. Patterns of linkage disequilibrium and long range hitchhiking in evolving experimental *Drosophila melanogaster* populations. Mol Biol Evol 32(2):495–509.

Garland M, Rose MR. 2009. Experimental evolution: concepts, methods, and applications of selection experiments. Berkeley, CA: University of California Press.

Hill WG, Robertson A. 1966. The effect of linkage on limits to artificial selection. Genet Res 8(3):269–294.

Huang Y, Wright SI, Agrawal AF. 2014. Genome-wide patterns of genetic variation within and among alternative selective regimes. PLoS Genet 10(8):e1004527.

Ignatiadis N, Klaus B, Zaugg J, Huber W. 2016. Data-driven hypothesis weighting increases detection power in genome-scale multiple testing. Nat Methods 13(7):577–580.

Kawecki TJ, Lenski RE, Ebert D, Hollis B, Olivieri I, Whitlock MC. 2012. Experimental Evolution. Trends Ecol Evol 27(10):547–560.

Kofler R, Schlötterer C. 2013. A guide for the design of evolve and resequencing studies. Mol Biol Evol 31(2):474–483.

Lenormand T, Chevin LM, Bataillon T. 2016. Parallel evolution: what does it (not) tell us and why is it (still) interesting? In: Ramsey G, Pence CH, editors. Chance in Evolution. Chicago, USA: Chicago University Press. p.196.

Lenski RE. 2017. Experimental evolution and the dynamics of adaptation and genome evolution in microbial populations. ISME J 11(10):2181–2194.

Long A, Liti G, Luptak A, Tenaillon O. 2015. Elucidating the molecular architecture of adaptation via evolve and resequence experiments. Nature Rev Genet 16(10):567–582.

Michalak P, Kang L, Scho MF, Garner H, Loeschcke V. 2019. Genomic signatures of experimental adaptive radiation in *Drosophila*. Mol Ecol 28(3):600–614.

Nouhaud P, Tobler R, Nolte V, Schlötterer C. 2016. Ancestral population reconstitution from isofemale lines as a tool for experimental evolution. Ecol Evol 6(20):7169–7175.

Nuzhdin SV, Turner TL. 2013. Promises and limitations of hitchhiking mapping. Curr Opin Genet Dev 23(6):694–699.

OrozcoterWengel P, Kapun M, Nolte V, Kofler R, Flatt T, Schlötterer C. 2012. Adaptation of *Drosophila* to a novel laboratory environment reveals temporally heterogenous trajectories of selected alleles. Mol Ecol 21(20):4931–4941.

Papkou A, Guzella T, Yang W, Koepper S, Pees B, Schalkowski R, Barg MC, Rosenstiel PC, Teotónio H, Schulenburg H. 2019. The genomic basis of Red Queen dynamics during rapid reciprocal host–pathogen coevolution. PNAS 116(3):923–928.

Phillips MA, Rutledge GA, Kezos JN, Greenspan ZS, Talbott A, Matty S, Arain H, Mueller L, Rose MR, Shahrestani P. 2018. Effects of evolutionary history on genome wide phenotypic convergence in Drosophila populations. BMC Genomics 116(1):743.

Plucain J, Suau A, Cruveiller S, Médigue C, Schneider D, Le Gac M. 2016. Contrasting the effects of historical contingency on phenotypic and genomic trajectories during a two-step evolution experiment with bacteria. BMC Evol Biol 16(1):86.

Porter ML, Crandall KA. 2003. Lost along the way: the significance of evolution in reverse. Trends Ecol Evol 18(10):541–547.

Remigi P, Masson-Boivin C, Rocha EPC. 2019. Experimental evolution as a tool to investigate natural processes and molecular functions. Trends Microbiol.

Schlötterer C, Kofler R, Versace E, Tobler R, Franssen S. 2015. Combining experimental evolution with next-generation sequencing: a powerful tool to study adaptation from standing genetic variation. Heredity 114(5):431–440.

Schlötterer C, Tobler R, Kofler R, Nolte V. 2014. Sequencing pools of individuals - mining genome-wide polymorphism data without big funding. Nat Rev Genet 15(11):749–763.

Schou MF, Kristensen TN, Kellermann V, Schlötterer C, Loeschcke C. 2014. A *Drosophila* laboratory evolution experiment points to low evolutionary potential under increased temperatures likely to be experienced in the future. J Evol Biol 27(9): 1859–1868.

Seabra SG, Fragata I, Antunes MA, Faria GS, Santos MA, Sousa VC, Simões P, Matos M. 2017. Different genomic changes underlie adaptive evolution in populations of contrasting history. MBE 35(3):549–563.

Simões P, Fragata I, Santos J, Santos MA, Santos M, Rose MR, Matos M, unpublished data, https://www.biorxiv.org/content/10.1101/579524v2, last accessed June 26, 2019. How phenotypic convergence arises in experimental evolution.

Spitzer K, Pelizzola M, Futschik A, unpublished data, https://arxiv.org/abs/1902.08127, last accessed June 26, 2019. Modifying the Chi-square and the CMH test for population genetic inference: adapting to over-dispersion.

Teotónio H, Chelo IM, Bradić M, Rose MR, Long AD. 2009. Experimental evolution reveals natural selection on standing genetic variation. Nat Genet 41(2):251–257.

Teotónio H, Rose MR. 2001. Perspectives: reverse evolution. Evolution 55(4):653–660.

ToblerR, Franssen SU, Kofler R, Orozco-terWengel P, Nolte V, Hermisson J, Schlötterer C. 2014. Massive habitat-specific genomic response in *D. melanogaster* populations during experimental evolution in hot and cold environments. Mol Biol Evol 31(2):364–375.

Turner T, Andrew S, Fields T, Rice WR, Tarone AM. 2011. Population-Based Resequencing of Experimentally Evolved Populations Reveals the Genetic Basis of Body Size Variation in *Drosophila melanogaster*. PLoS Genet 7(3):e1001336.

Turner T, Miller P. 2012. Investigating Natural Variation in Drosophila Courtship Song by the Evolve and Resequence Approach. Genetics 191(2):633–642.

