## Supplement Material for "Secondary evolve and re-sequencing: an experimental confirmation of putative selection targets without phenotyping"

FIGURES LEGENDS

**Figure SI 1** Close up of the two focal candidate sets: $FET_x$ (left) and $CMH_{y,z}$ (right).

The region of +/- 300kbp around the most significant SNP from the $CMH_{x,y,z}$ and $FET_x$ is indicated by gray and brown segments, which. The 20 most significant SNPs from $FET_x$ and $CMH_{y,z}$ are shown in red and blue.

**Figure SI 2** Allele frequency spectrum of all candidate SNPs from the primary E&R (red) relative to background SNPs (black) on intersected 3R arm region in the starting populations. Each panel shows the fraction of SNPs (y-axis) in allele frequencies bins (x-axis) for experiment x (top panels) and z (bottom panels). Frequencies bin sizes 0.02.

**Figure SI 3** Allele frequency changes of the 20 most significant SNPs from $FET_x$ (red) and $CMH_{y,z}$ (blue) for the primary E&R (generation F0-F70) and secondary E&R (D0-D30).

Each panel shows another replicate population from the secondary E&R. Top row experiment x, bottom row experiment z.

**Figure SI 4** Repeatability of selection signatures in primary and secondary E&R.

Corresponds to figure 4, but data are shown for all replicates of the secondary E&R experiments.

**Figure SI 5** Repeatability of selection signatures in primary and secondary E&R.

Corresponds to figure 4, but data are shown for all replicates of the secondary E&R experiments.

**Figure SI 6** Close up of the Manhattan plots in the secondary E&R study showing the p-values for the CMH from experiment x (left) and z (right). See Fig. SI 1 for more details.

TABLES

**Table SI 1** Sequencing information.

| **Library** | Primary E&R experiment: Ancestral replicates (F0) | Primary E&R experiment: Evolved replicates (F70) and Secondary E&R experiment: Diluted ancestral (D0) and evolved replicates (D30) |
|---|---|---|
| **Method for DNA isolation** | High salt extraction protocol including RNase A treatment from Miller et al. 1988 | High salt extraction protocol including RNase A treatment from Miller et al. 1988. |
| **Amount of starting material (genomic DNA)** | 5μg | 1μg |
| **Fragmentation method** | Covaris S2[1] | Covaris S2[1] |
| **Kit** | TruSeq DNA PCR-Free Sample Prep Kit[2] | NEBNext Ultra DNA II Library Prep Kit[3] |
| **Size selection method** | AMPure XP beads[4] | AMPure XP beads[4] |
| **Insert size (bp)** | 380 | 260 |
| **Polymerase** | PCR-free | Q5 Mastermix |
| **No. of PCR cycles** | PCR-free | 4 |
| **Sequencing platform** | HiSeq 2000 | HiSeq XTEN |
| **Read length (bp)** | 2x100 | 2x150 |
| **SRA Run accession no.** | XXX | XXX |

**1) Covaris, Inc. Woburn, MA, USA**

**2) Illumina, San Diego, CA**

**3) New England Biolabs, Ipswich, MA**

**4) Beckman Coulter, Carlsbad, CA**

**Table SI 2** Candidates list from the primary E&R study.

| | raw number of variants | | | filtered number of variants | | | intersection over the 3 mappers | coverage filtering and mapping outliers filters |
|---|---|---|---|---|---|---|---|---|
| | NovoAlign | Bowtie2 | BWA-MEM | NovoAlign | Bowtie2 | BWA-MEM | | |
| SNPs autosomes/ X | 3988599/ 636251 | 3905323/ 622883 | 4010418/ 640279 | 2865514/ 452947 | 2737761/ 431690 | 2876308/ 454144 | 2532855/ 404290 | 2225925/ 334613 |
| INDELs autosomes/ X | 404255/ 85078 | 314352/ 69037 | 394719/ 83311 | | | | | |

**Table SI 3** Average coverage from sync file at SNP positions per sample for NovoAlign counts for primary (secondary) E&R.

| | Primary E&R | | Secondary E&R | |
|---|---|---|---|---|
| | F0<br>autosomes / X | F70<br>autosomes / X | D0<br>autosomes / X | D30<br>autosomes / X |
| **Replicate x**<br>x.1-x.2 | 167/167 | 89/77 | 54-49/47-42 | 67-67/59-56 |
| **Replicate y** | 256/256 | 79/68 | - | - |
| **Replicate z**<br>z.1-z.2-z.3 | 185/185 | 88/74 | 58-80-37/50-66-32 | 71-72-78/59-60-65 |

FIGURES

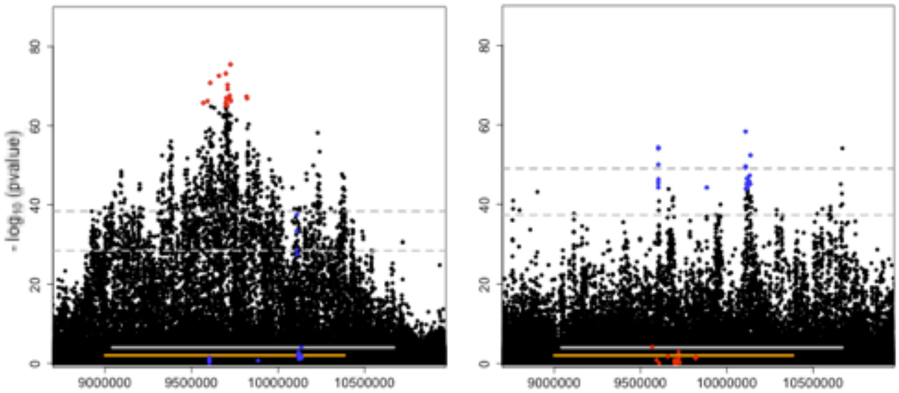


**Figure SI 1** Close up of the two focal candidate sets: $FET_x$ (left) and $CMH_{y,z}$ (right).

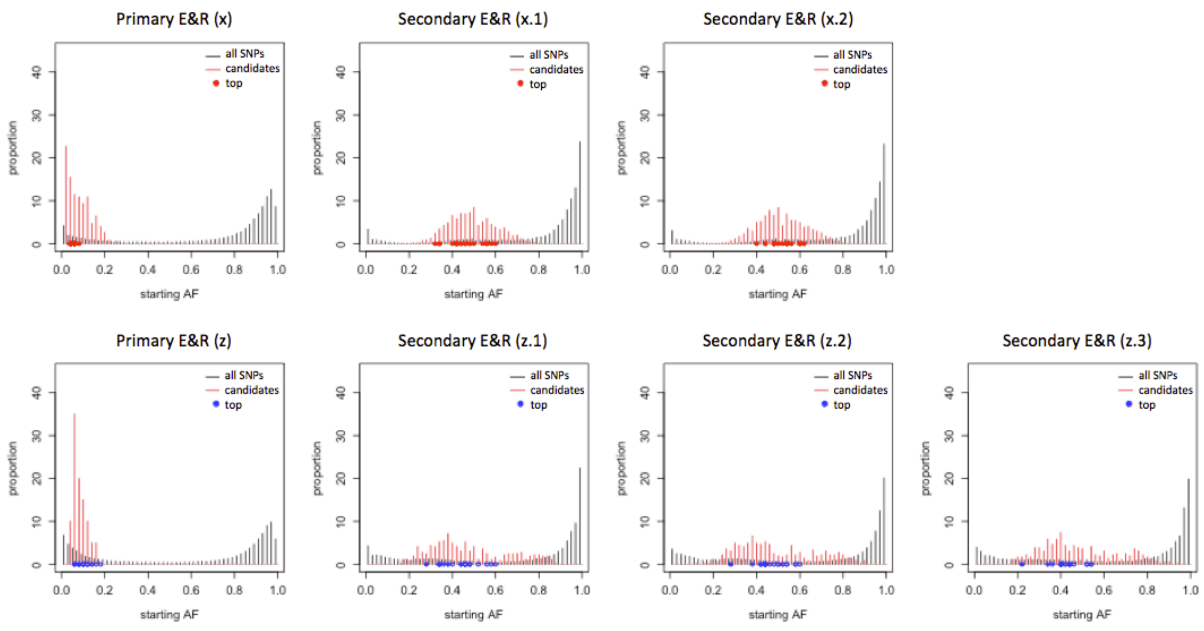


**Figure SI 2** Allele frequency spectrum of all candidate SNPs from the primary E&R (red) relative to background SNPs (black) on intersected 3R arm region in the starting populations.

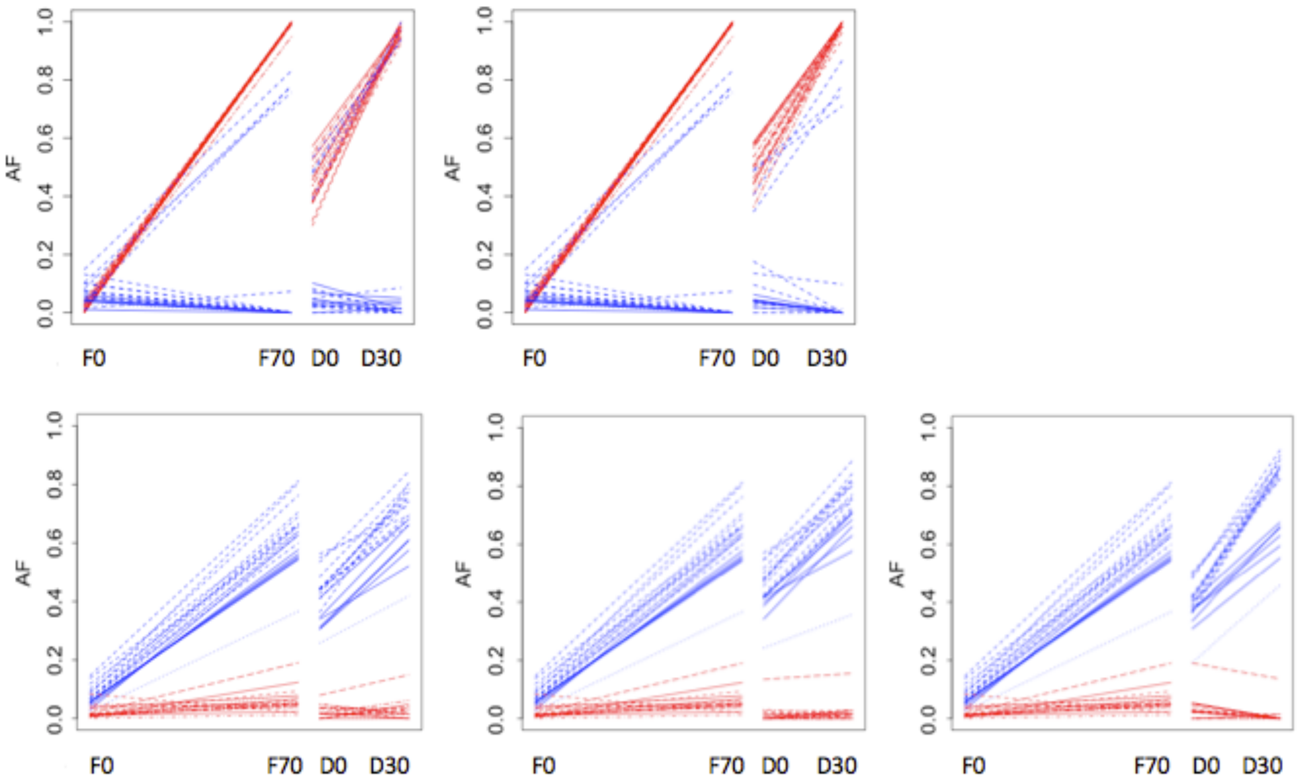


**Figure SI 3** Allele frequency changes of the 20 most significant SNPs from $FET_x$ (red) and $CMH_{y,z}$ (blue) for the primary E&R (generation F0-F70) and secondary E&R (D0-D30).

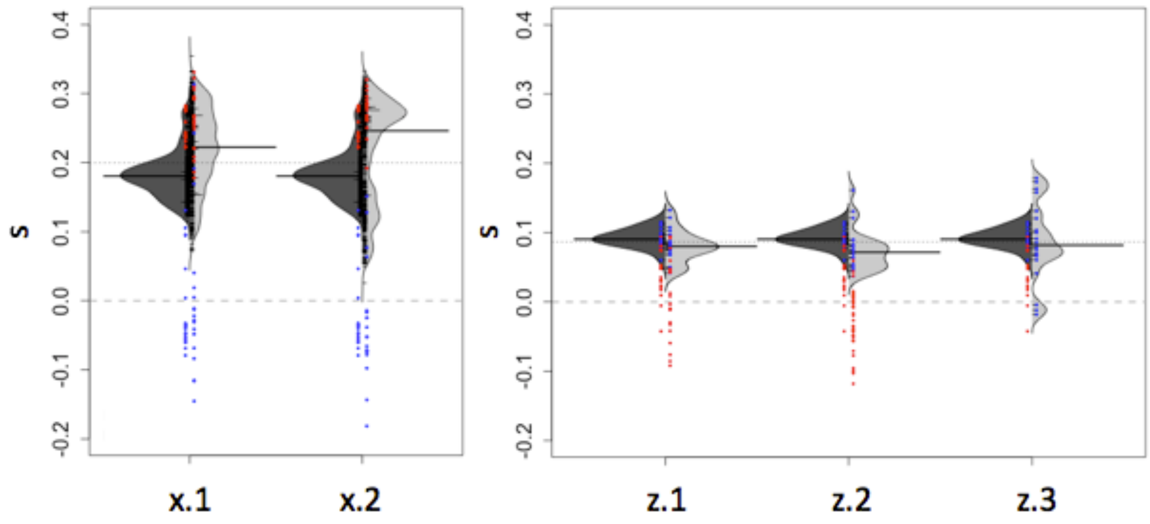


**Figure SI 4** Repeatability of selection signatures in primary and secondary E&R.

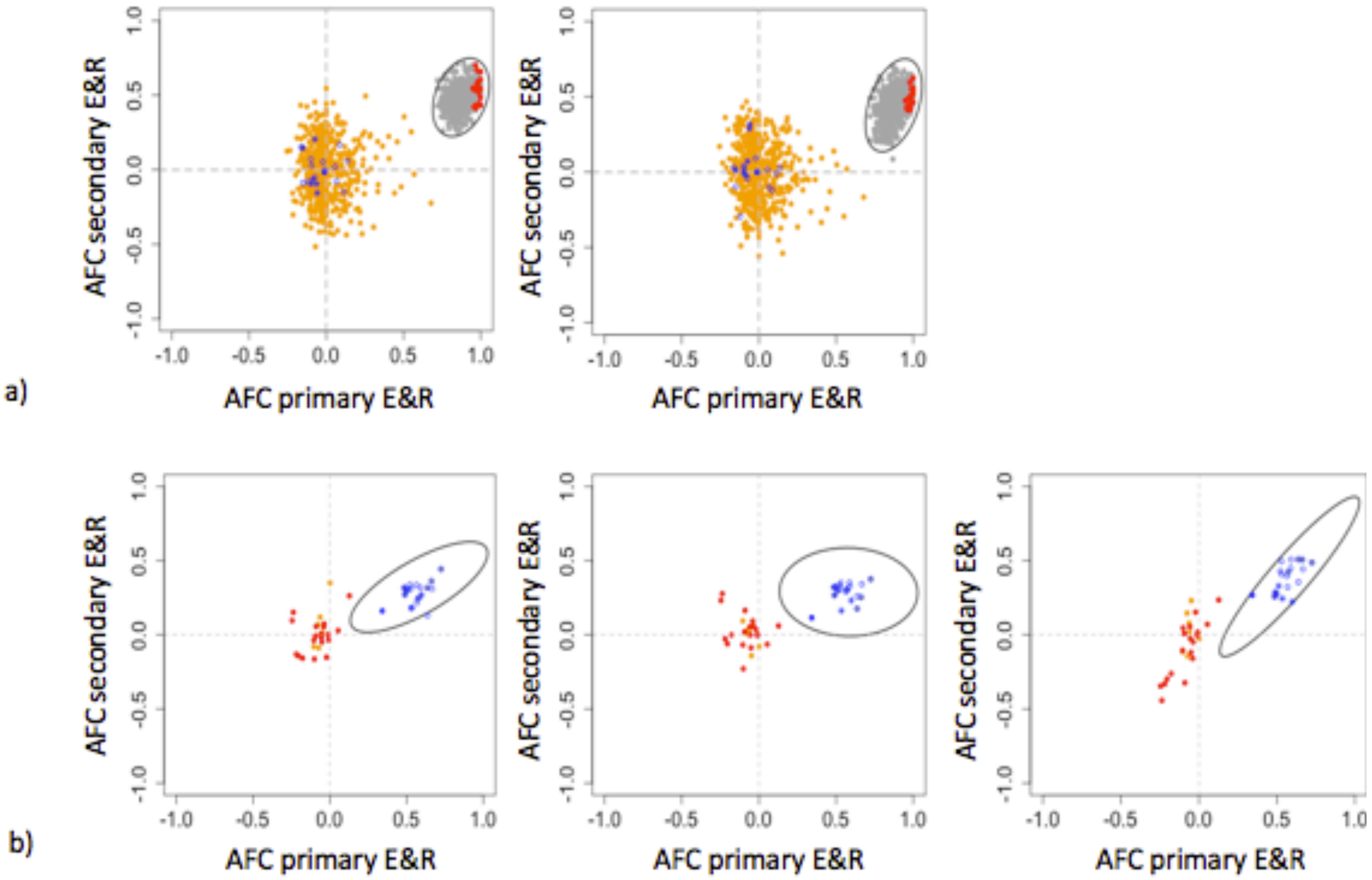


**Figure SI 5** Repeatability of selection signatures in primary and secondary E&R.

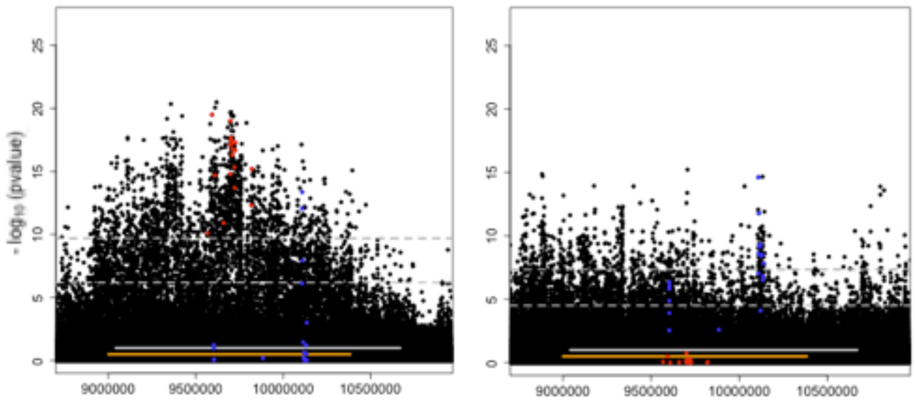


**Figure SI 6** Close up of the Manhattan plots in the secondary E&R study showing the p-values for the CMH from experiment x (left) and z (right).

SUPPLEMENTARY MATERIAL

**The primary E&R experiment**

**Creation of a Bona Fide SNP Catalogue for the Primary E&R study**

Quality Control. FASTQ files were first demultiplexed using ReadTools (Gómez-Sánchez and Schlötterer 2018) (version 1.3.0; AssignReadGroupByBarcode --splitSample, --maximumMismatches 1, providing the corresponding barcodes). The raw reads were assessed for their quality using the FastQC software (http://www.bioinformatics.babraham.ac.uk/projects/fastqc/). Low quality bases at the 3′-end of each read were trimmed using ReadTools (--mottQualityThreshold 18, --minReadLength 50, --disable5pTrim true) and the resulting BAM files were converted to compressed FASTQ files using ReadTools (ReadsToFastq --interleavedInput true --barcodeInReadName true --outputFormat GZIP).

Mapping. Processed paired-end reads were mapped to the indexed *D. simulans* reference genome (Palmieri et al. 2015) using i) NovoAlign (http://novocraft.com) (version 3.03.02; -i 250,100 -F STDFQ -o SAM -r RANDOM), ii) Bowtie2 (Langmead and Salzberg 2012) (version 2.2.6; -q --phred33 --end-to-end -X 1500) and iii) BWA-MEM (Li and Durbin 2009) (version 0.7.13; bwamem) on a Hadoop cluster with DistMap (Pandey and Schlötterer 2013) (version 2.7.5). This procedure generated three BAM files per sample.

Mapping filtering. PCR duplicates were removed using the PICARD MarkDuplicates tool (version 2.1.0; REMOVE_DUPLICATES=true VALIDATION_STRINGENCY=SILENT) (http://broadinstitute.github.io/picard/). We kept mapped reads with each segment properly aligned and removed reads that mapped equally well to multiple positions or had a low mapping quality using SAMtools (Li et al. 2009) (version 1.1; -b -q 20 -f 0x002 -F 0x004 -F 0x008). Files were again sorted and indexed via SAMtools. Summaries of mapped reads were obtained with SAMtools (flagstat) and empirical average coverages were reported (Supplementary Table II).

SNPs calling. SNPs were called separately for each mapper using Freebayes (Garrison and Marth, unpublished data) with freebayes-parallel (version 1.2.0-2-g29c4002; --min-alternate-count 2, --min-coverage 5, --min-alternate-fraction 0.01, --min-base-quality 40 --pooled-continuous) from a merged BAM file of all processed F0 and F70 replicate pools. The parallelization required a partitioning of the reference genome which was achieved by providing a so-called region file. We chose to split the genome at sites with TE insertions, and generated the corresponding region file from the annotation of repeats used by Barghi et al.

2019 after having combined overlapping repeated regions using BEDTools (Quinlan and Hall 2010; version 2.27.1; sort, merge, sort successively). The reference cutting points were determined as the boundaries of the overlapping regions, conditioned on a >200 bp-length, using a custom R script.

SNPs filtering. We obtained one list of variants per in a VCF format (Danecek et al. 2011). To obtain a final set of SNPs, each raw list of variants required to be filtered

i) to remove variants based on depth (i.e. coverage) at the variant position using BCFtools (filter –i SAF>0 && SAR>0 && SRF>0 && SRR>0 && RPR>0 && RPL>0 && (SAF+SAR+SRF+SRR)>4 && (SAF+SAR+SRF+SRR)<threshold). The threshold corresponds to the 99% quantile of high quality base reads number over all samples at the variant position, for each mapper and for X chromosome and autosomes separately (NovoAlign: 641/653, Bowtie2: 529/562, BWA-MEM 638/650). These coverage values were obtained after having converted the raw VCF to a readable text file using BCFtools (query -f '%CHROM %POS %SAF %SAR %SRF %SRR %DP\n' raw_map_chr.vcf | awk '{printf "%s %s %s %s %s %s %s %i %f\n",$1,$2,$3,$4,$5,$6,$7,$3+$4+$5+$6,($3+$4)/($3+$4+$5+$6)}' | sed '1 i\ CHROM POS SAF SAR SRF SRR DP DPQ AF|' | tr '|' '\n' > raw_chr_map.txt).

ii) to remove SNPs within 5bp of an INDEL using BCFtools (filter -g 5),

iii) to keep SNPs only using BCFtools (filter -i 'TYPE="snp"'),

iv) to keep SNPs with at most two alleles denoted as reference and alternate alleles with VCFtools (Danecek et al. 2011; version 0.1.15; --vcf --min-alleles 2 --max-alleles 2 --recode-INFO-all --stdout --recode),

v) to simplify multi-nucleotide polymorphisms into SNPs using vt (Tan et al. 2015; decompose_blocksub, normalize successively). vt was cloned from the github repository (https://github.com/atks/vt) in December 2018 (version 0.57721).

vi) to mask SNPs found in repeated regions using BEDtools (intersect -v -wa –header).

VCF files were first bgzipped and tabix indexed (Li 2011; version 1.8; -p vcf), and then intersected between the 3 mappers to discard SNPs that differ in identity between mappers using VCFtools (vcf-isec –n=3).

This procedure obtained an intermediate list of 2,937,145 SNPs. The number of raw and filtered variants identified is listed in Supplementary Table II obtained with BCFtools (version 1.8; stats).

Triple mapping procedure. Processed BAM files were first converted in pileup format using SAMtools (mpileup –B, -Q0, -d10000) and then to synchronized – sync - files using the popoolation2 (Kofler et al. 2011) tool mpileup2sync.jar (version 1201; --min-qual 20 --fastq-type sanger). sync files were subset at the SNPs positions using a custom R script. For each mapper, SNPs with extremely low or high coverage in a least one sample (< 1st quantile or > 99th quantile) were removed. SNPs with the same frequency in all samples were removed. This procedure ensured to only use reliable positions with a base quality of 20 to compute the count and frequency of each SNP. The minor allele was assessed using the poolSeq::read.sync R function (Taus et al. 2017; version 0.3.2; polarization=”minor”). We then performed Fisher Exact Tests within the replicates of F0 and within F70 (2 rows) between the mappers (3 columns) and filtered against SNPs that significantly differed in allele frequency at a 1 % FDR. This leads to a final set of 2,560,538 SNPs.

**Identifying Regions under Selection in the primary E&R**

The frequencies and counts of the rising allele in the corresponding replicate were calculated using the poolSeq::read.sync R function (polarization=”rising”) from Novoalign.

Estimation of $N_e$. From 100 trials of random subsets of 1,000 SNPs, per replicate and for X chromosose and autosomes separately, $N_e$ was estimated using the median over all trials from the poolSeq::estimateNe R function (method=”P.planI”, truncAF=0.01, Ncensus = 1,250, all other arguments set to default) (Jónás et al. 2016; Taus et al. 2017).

Neutral Simulations. To determine significance cut offs we ran neutral forward Wright-Fisher simulations based on the determined $N_e$ estimates per replicate providing the empirical starting allele frequencies using first poolSeq::wf.traj and then poolSeq::sample.alleles (coverage mode) R functions to incorporate sampling noise. The same FET and CMH test are applied to the simulated data to identify candidates separately for each replicate(s).

FET and CMH tests p-values weighted by the Iterative Hypothesis Weighting procedure (IHW). To contrast ancestral and evolved populations in the primary E&R study, Cochran-Mantel-Haenszel (CMH) (Agresti 2002) were done using 3 (x, y, z) or 2 (y, z) replicates. For the comparison of the two time points in single replicates, Fisher´s exact tests (FET) were used. CMH tests were done using the poolSeq::cmh.test R function (min.cov=1, max.cov=1, min.cnt=1, all other arguments set to default) and FET were done using the fisher.test R function. We identified candidate SNPs from the primary E&R using the IHW procedure. To this end we performed neutral forward Wright-Fisher simulations without linkage based on the $N_e$ estimates for each replicate population and the empirical starting allele frequencies using poolSeq::wf.traj. We incorporated in a second step sampling noise using poolSeq::sample.alleles (coverage mode). Both CMH tests and FET were done on the neutral simulated data. The IHW procedure (Ignatiadis et al. 2016) was applied separately for both observed and simulated counts using the mean coverage per SNP as a covariate, using the IHW::ihw (version 1.8.0) and IHW::adj_pvalues R functions; for a FET, this would be the average coverage over two time points of one replicate whereas for a CMH test, this would be the average coverage over >1 replicates and two time points.

Estimation of selection coefficients *s*. Selection coefficients were estimated per SNP per population using the poolSeq::estimateSH R function, with the default dominance coefficient of 0.5, on pseudo-count data to avoid any SNPs reaching an absorbing state (i.e. fixation or loss) not having an *s* estimate. Per SNP, we rescaled the count of the rising allele by adding (subtracting) 1 to the total depth if the SNP got lost (fixed) at any time point within any replicate.

### REFERENCES


Agresti A. 2002. Categorical Data Analysis. John Wiley & Sons, 2nd Edition.

Barghi N, Tobler R, Nolte V, Jakšić AM, Mallard F, Otte KA, Dolezal M, Taus T, Kofler R, Schlötterer C. 2019. Genetic redundancy fuels polygenic adaptation in *Drosophila*. *PLoS Biol* 17(2):e3000128.

Cleary JG, Braithwaite R, Gaastra K, Hilbush BS, Inglis S, Irvine SA, Jackson A, Littin R, Nohzadeh-Malakshah S, Rathod M, et al. 2014. Joint variant and de novo mutation identification on pedigrees from high-throughput sequencing data. *J Comput Biol* 21(6):405–419.

Danecek P, Auton A, Abecasis G, Albers CA, Banks E, DePristo MA, Handsaker RE, Lunter G, Marth GT, Sherry ST, et al. 2011. The variant call format and VCFtools. *Bioinformatics* 27(15):2156–2158.

Li H, Handsaker B, Wysoker A, Fennell T, Ruan J, Homer N, Marth G, Abecasis G, Durbin R. 2009. The Sequence Alignment/Map format and SAMtools. *Bioinformatics* 25(16):2078-2079.

Garrison E, Marth G, unpublished data, https://arxiv.org/abs/1207.3907, last accessed June 26, 2019. Haplotype-based variant detection from short-read sequencing.

Gómez-Sánchez D, Schlötterer C. 2018. ReadTools: a universal toolkit for handling sequence data from different sequencing platforms. *Mol Ecol Resour* 18(3):676-680.

Kofler R, Langmüller AM, Nouhaud P, Otte KA, Schlötterer C. 2016. Suitability of different mapping algorithms for genome-wide polymorphism scans with pool-seq data. *G3-Genes Genom Genet* 6(11):3507-3515.

Kofler R, Pandey RV, Schlötterer C. 2011. PoPoolation2: identifying differentiation between populations using sequencing of pooled DNA samples (Pool-Seq). *Bioinformatics* 27(24):3435-3436.

Quinlan QR, Hall IM. 2010. BEDTools: a flexible suite of utilities for comparing genomic features. *Bioinformatics* 26(6):841-842.

Ignatiadis N, Klaus B, Zaugg J, Huber W. 2016. Data-driven hypothesis weighting increases detection power in genome-scale multiple testing. *Nat Methods* 13(7):577-580.

Jónás Á, Taus T, Kosiol C, Schlötterer C, Futschik A. 2016. Estimating the effective population size from temporal allele frequency changes in experimental evolution. *Genetics* 204(2):723-735.

Langmead B, Salzberg SL. 2012. Fast gapped-read alignment with Bowtie2. *Nat Methods* 9(4):357-359.

Li H. 2011. Tabix; fast retrieval of sequence features from generic TAB-delimited files. *Bioinformatics* 27(5):718-719.

Li H, Durbin R. 2009. Fast and accurate short read alignment with Burrows-Wheeler transform. *Bioinformatics* 25(14):1754-1760.

Li H, Handsaker B, Wysoger A, Fennell T, Ruan J, Homer N, Marth G, Abecasis G, Pandey RV, Schlötterer C. 2013. DistMap: a toolkit for distributed short read mapping on a Hadoop cluster. *PLoS One* 8(8):e72614.

Miller S, Dykes DD, Polesky HF. 1988. A simple salting out procedure for extracting DNA from human nucleated cells. *NAR* 16(3):1215.

Nouhaud P, Tobler R, Nolte V, Schlötterer C. 2016. Ancestral population reconstitution from isofemale lines as a tool for experimental evolution. *Ecol Evol* 6(20):7169-7175.

Palmieri N, Nolte V, Chen J, Schlötterer C. 2015. Genome assembly and annotation of *Drosophila simulans* strain from Madagascar. *Mol Ecol Resour* 15(2):372-381.

Strimmer K. 2008. A unified approach to false discovery rate estimation. *BMC Bioinformatics* 9(1):303.

Tan A, Abecasis GR, Kang HM. 2015. Unified representation of genetic variants. *Bioinformatics* 31(13):2202-2204.

Taus T, Futschik A, Schlötterer C. 2017. Quantifying selection with pool-seq time series data. *Mol Biol Evol* 34(11):3023-3034.

Vlachos C, Burny C, Pelizzola M, Borges R, Futschik A, Kofler R, Schlötterer C, unpublished data, https://www.biorxiv.org/content/10.1101/641852v1.abstract, last accessed June 26, 2019. Benchmarking software tools for detecting and quantifying selection in Evolve and Resequencing studies.
